# The Formation of Biogenic Guanine Crystals Closely Resembles Melanosome Morphogenesis

**DOI:** 10.1101/2022.10.01.510454

**Authors:** Avital Wagner, Alexander Upcher, Raquel Maria, Thorolf Magnesen, Einat Zelinger, Graça Raposo, Benjamin A. Palmer

**Affiliations:** Department of Chemistry, Ben-Gurion University of the Negev, Beer-Sheba 8410501, Israel; Ilse Katz Institute for Nanoscale Science & Technology, Ben-Gurion University of the Negev, Beer-Sheba 8410501, Israel; Department of Biological Sciences, University of Bergen, Postbox 7803, Bergen N-5020; The CSI Center for Scientific Imaging, The Robert H. Smith Faculty of Agriculture, Food and Environment, The Hebrew University of Jerusalem, POB 12, Rehovot 7610001, Israel; Institut Curie, PSL Research University, CNRS, UMR144, Structure and Membrane Compartments, 75005 Paris, France; Institut Curie, PSL Research University, CNRS, UMR144, Cell and Tissue Imaging Facility (PICT-IBiSA), 75005 Paris, France

## Abstract

Animals precisely control the morphology and assembly of highly reflective guanine crystals to produce diverse optical phenomena used in coloration and vision. However, little is known about how organisms regulate crystallization to produce optically useful crystal morphologies which express highly reflective crystal faces. Guanine crystals form inside iridosome vesicles within specialized chromatophore cells called iridophores. By following the formation of iridosomes in developing scallop eyes we show that, despite their different physical properties, the morphogenesis of iridosomes bears a striking resemblance to melanosome morphogenesis in vertebrates. A key finding is that pre-assembled intravesicular fibrillar sheets template crystal nucleation, growth, and orientation, dictating the formation of optically functional, but thermodynamically disfavored plate-like habits. The common control mechanisms for melanin and guanine formation inspire new approaches for manipulating the morphologies and properties of molecular materials.

## Introduction

Understanding how organisms control the formation of biogenic crystals is the central, widely studied question of (inorganic) biomineralization^1–6^. However, the mechanisms underlying the formation of organic bio-crystals, which are ubiquitous in animals^6–9^ and plants^10^, remain a mystery. Guanine crystals are the most widespread molecular bio-crystal. The crystals are constructed from π-stacked, H-bonded layers^11^, and exhibit exceptional optical properties, due to the extreme in-plane refractive index (n = 1.83) (Supplementary Fig. 1). By precisely controlling the shape and assembly of these crystals, organisms produce an array of different optical phenomena, used in camouflage^12^, display^13^ and vision^14,15^. From a single material, an impressive variety of crystal morphologies are generated including regular squares and hexagons, irregular polygons, and prisms^16^. All these crystals share a common feature – they preferentially express the highly reflective (100) face (parallel to the high-index, H-bonded layers) to maximize reflection. In contrast, guanine crystals grown *in vitro* from aqueous solutions grow along the thermodynamically favored, π-stacking direction resulting in the expression of low-index (∼1.45) crystal faces. It is not known how organisms exquisitely regulate crystallization to generate optically useful morphologies, but recent studies have ruled out several hypotheses: (i) the use of purines as intracrystalline growth additives^16^, (ii) molding of amorphous precursors^17^, and (iii) physical confinement by membranes^17^.

The little we know about guanine formation is that the crystals form inside membrane-bound vesicles called iridosomes^18,19^ within specialized chromatophore cells called iridophores. In vertebrates, iridophores differentiate from the same progenitor cell as melanophores (melanin pigment cells) and xanthophores (pteridine/carotenoid pigment cells)^20^. Bagnara et al., postulated that all vertebrate pigment organelles (‘-somes’) derive from a common ‘primordial vesicle’^21^, originating from the endosomal pathway^19^. Furthermore, it was shown that many of these pigment granules are lysosome-related organelles (LRO)^22-24^. Despite their inclusion in the LRO family, iridosomes differ from other pigments in that they generate structural colors from crystals, rather than absorptive colors from pigments. In contrast to other pigment organelles^19^ almost nothing is known about iridosome biogenesis. By following the formation of guanine crystals in a developing organism, we reveal key features of iridosome morphogenesis and show that it is strikingly similar to melanosome formation in vertebrates. Elucidating this pathway provides an answer to how organisms control guanine crystal morphology *via* the use of templating macromolecules.

## Results

The image forming mirror of scallop eyes illustrates the precise control which organisms exert over crystal morphology and assembly. Adult *Pecten maximus* scallops have hundreds of eyes (Fig. 1A), each containing a concave mirror (Fig. 1B, pseudo colored green) which focuses light onto the overlying retinas^25^. The mirror, which is contained in an iridophore cell (Supplementary Fig. 2), is formed from tessellated layers of square guanine crystals (Fig. 1C-D) – a crystal habit forbidden by the monoclinic symmetry of guanine^11,26^. Each crystal is tightly bound by an enveloping membrane (i.e., an iridosome organelle)^25^ (insert; Fig. 1C). The highly reflective (100) face of the crystals is oriented towards the incident light across the entire surface of the mirror to direct light to focal points on the two retinas^14,25^.

**Figure 1.**
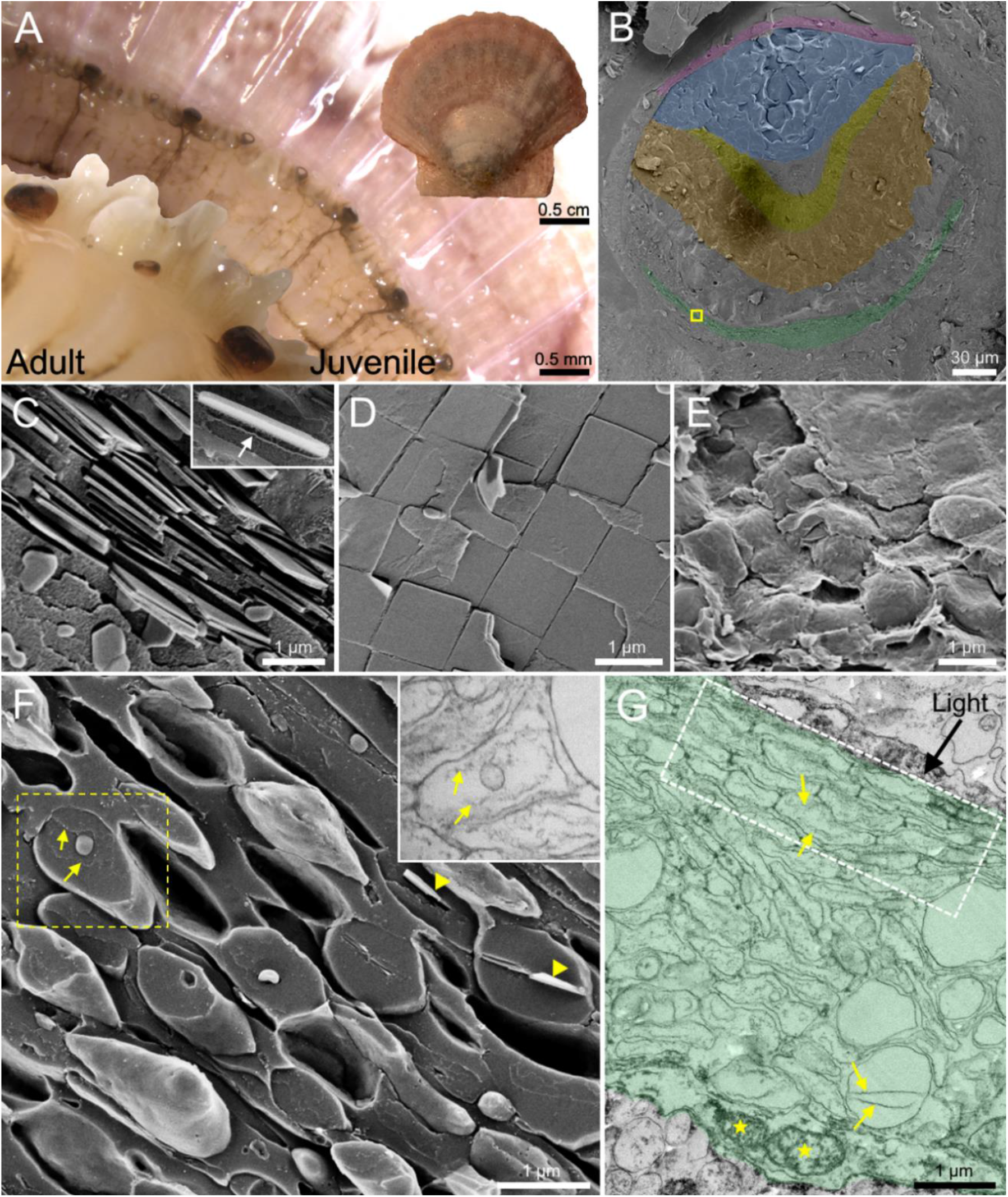
The image-forming mirror in the eyes of adult and juvenile scallops. **(A)** Stereomicroscope images of the upper mantle in an adult and juvenile scallop, containing numerous eyes. Insert; a juvenile scallop (20 mm in diameter). **(B)** Pseudo-colored cryo-SEM micrograph of a freeze-fractured eye of a juvenile scallop approximately 230 μm in diameter. Cornea; purple, blue; lens, yellow; distal retina, orange; proximal retina, green; the mirror region. **(C)-(D)** Cryo-SEM micrographs of the mirror region in an adult scallop viewed perpendicular to (C) and along (D) the optic axis. Insert in (C) shows the delimiting membrane of the iridosome (white arrow) around a fully formed crystal. **(E)-(F)** Cryo-SEM micrographs (from the yellow region of interest in (B)) of the mirror in a juvenile scallop eye viewed along (E) and perpendicular (F) to the optic axis. The mirror is composed of ellipsoidal iridosomes at various formation stages. Insert in (F); SEM image of an iridosome taken with STEM detector showing an example of a vesicle containing two intraluminal fibrils. **(G)** SEM image taken with STEM detector of an ultra-thin tissue section, showing the mirror in the juvenile scallop eye. In (F) and (G) a variety of iridosome formation states are observed. (F),(G): yellow arrows; intraluminal fibrils, yellow stars; mitochondria, white dashed box; an area of tessellated vesicles.

To investigate the formation of the unusual guanine crystals, we examined juvenile (∼two months post hatching) scallop eyes (Fig. 1A-B) using cryogenic scanning (cryo-SEM) and transmission electron microscopy (TEM). The 10-20 mm scallops (Fig. 1A), have dozens of minute eyes, 150 to 250 μm in diameter (Fig. 1B). In juveniles, the mirror is composed of loosely tessellated, ‘pillow-shaped’ iridosomes (Fig. 1E) rather than the perfectly tiled squares of the adult (Fig. 1D). Cryo-SEM images of longitudinal sections, show that the mirror is filled with spherical and ellipsoidal iridosomes that, when elongated, have their long axis oriented perpendicular to the incident light (Fig. 1F-G). A variety of morphologically distinct iridosomes are observed in the forming iridophore cell, representing different stages of iridosome development (Fig. 1F-G). Mature iridosomes, typically found at the distal edge of the cell (towards the incident light direction), are elongated, contain partially formed crystals (Fig. 1F yellow arrowheads) and display the characteristic tessellated suprastructure of adults (Fig. 1E-G). Immature iridosomes, located proximally in the cell are more spherical and lack crystals (Fig. 1F-G). Prevalent features of these immature iridosomes are two intraluminal fibrils (Fig. 1F-G, yellow arrows) and intraluminal vesicles (ILVs) of various sizes (Fig. 1F-G). The distinct iridosome morphological states together with a proposed morphogenesis, derived by following guanine nucleation and growth, are shown in Fig. 2.

**Figure 2.**
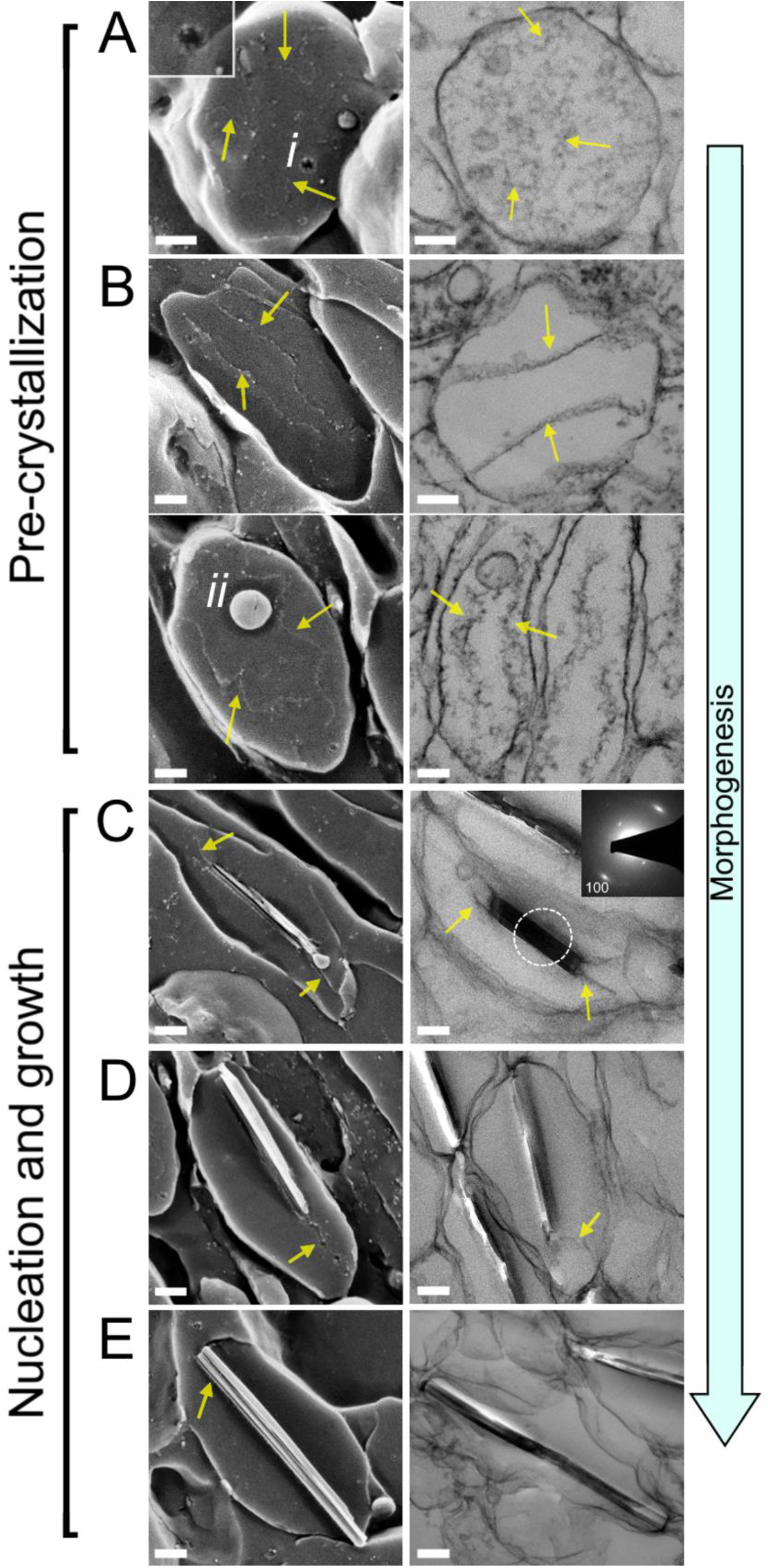
The morphogenesis of guanine crystals in juvenile scallop eyes by cryo-SEM (left column) and TEM (right column). **(A)-(C)** Pre-crystallization: **(A)** Spherical vesicles containing small type *i* intraluminal vesicles and some disordered fibrils. Insert; high magnification image of a type *i* intraluminal vesicle. **(B)** Ellipsoidal iridosomes characterized by two intraluminal fibrils stretching across the length of the vesicle. Ellipsoidal iridosomes prior to crystallization are characterized by two types of intraluminal vesicles. Type *ii* vesicles shown here, are bound to the intraluminal fibers. **(C)-(E)** Nucleation and growth: **(C)** As morphogenesis proceeds, the iridosomes elongate and a small guanine crystal nucleates between the fibrils. The thin immature crystals are elongated along the H-bonded direction, and exhibit an electron diffraction pattern characteristic of *β-*guanine (insert). **(D)** The intraluminal fibrils template the growth of the crystal along the H-bonded direction. The fibrils become tightly bound to the (100) face of the crystals. **(E)** Mature crystals stretch and re-shape the iridosome membrane, which condenses on the surface of the faceted crystal. Scale bars; 100 nm, yellow arrows; intraluminal fibrils.

The proximal region of the iridophore cell contains numerous spherical vesicles ∼500 nm in diameter, which contain multiple smaller (25-50 nm) intraluminal vesicles (type *i*) and some disordered fibrils (Fig. 2A). These vesicles closely resemble ‘premelanosomes’ – multi-vesicular bodies (MVBs) characterizing the first stage of melanosome organellogenesis^27^. We thus assign these vesicles to the most premature iridosomes. Type *i* intraluminal vesicles, which have a rough surface texture (Fig. 2A insert and Supplementary Fig. 3), are observed only in spherical iridosomes, associated with the earliest formation stages. During the next formation stage, the iridosomes elongate concomitantly with the formation of two intraluminal fibrils which stretch from one pole of the vesicle to the other (Fig. 2B)^28^. Two types of intraluminal vesicles are seen in these iridosomes. Type *ii* vesicles (80-100 nm) are bound to one of the fibrils, and type *iii* vesicles (250-275 nm) are typically unbound (Supplementary Fig. 3).

As morphogenesis proceeds, a small guanine crystal nucleates between the two fibrils (Fig. 2C). The crystallinity was confirmed by *in situ* electron diffraction which exhibits the characteristic signature of *β*-guanine. The guanine crystal then grows across the iridosome along the H-bonding direction, tracking along the fibrils which become tightly bound to the (100) crystal faces (Fig. 2E-F). Eventually, the growing crystal contacts with both sides of the iridosome, pulling the membrane around the growth front of the crystal. The re-shaping of the iridosome vesicle by the growing crystal was also observed in spiders^17^ and lizards^29^, and again shows that crystal habit is not dictated by physical confinement by the vesicle membrane.

In adult scallops, the iridosome membrane and intraluminal fibrils ultimately become fused to the crystal surface (Fig. 3, Fig. 1C, insert), assuming its faceted shape^14^. This two-tiered membrane/fibril crystal envelope provides an explanation to previous reports of ‘double-membrane’ delimited iridosomes (Supplementary Fig. 4)^18^.

**Figure 3.**
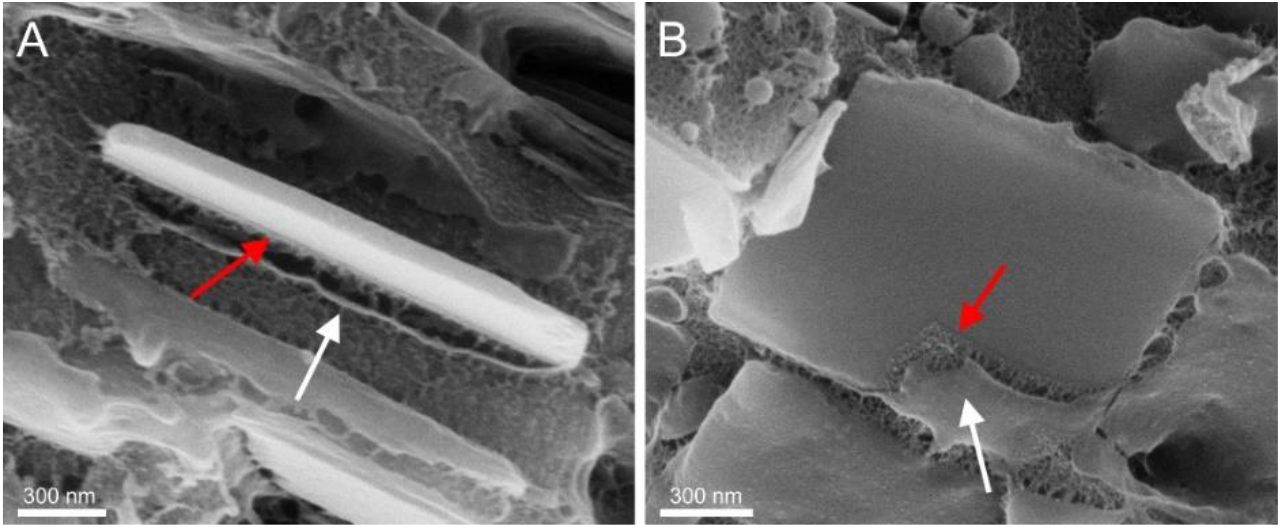
Cryo-SEM images of guanine crystals from an adult scallop eye. **(A)** side view **(B)** top-down view of guanine crystals showing both the outer lipid vesicle membrane (white arrows) and an inner layer tightly bound to the crystal (red arrows). We assign the inner enveloping layer to the fibrillar sheets observed in juvenile scallop iridosomes which become fused to the crystal surface.

To further investigate the nature and function of the intraluminal fibrils, we performed TEM tomography on iridosomes from juvenile scallops. Three-dimensional tomographic reconstructions of early iridosomes revealed that the two intraluminal ‘fibrils’ are in fact 2D sheets (Fig. 4A-D, Supplementary movies 1-2). These sheets act as a template for guanine nucleation and ‘cap’ the (100) faces of the growing crystal. Macromolecular templating effectively forces the crystals to grow along H-bonding direction resulting in the expression of the highly reflective (100) face. This finding reveals how organisms inhibit crystal growth along the thermodynamically favored π-stacking growth direction to preferentially form optically functional (100) plates. Observations of intercalating sheets in guanine crystals in spiders^17^ suggests that templated growth could be a general feature of biogenic guanine formation.

**Figure 4.**
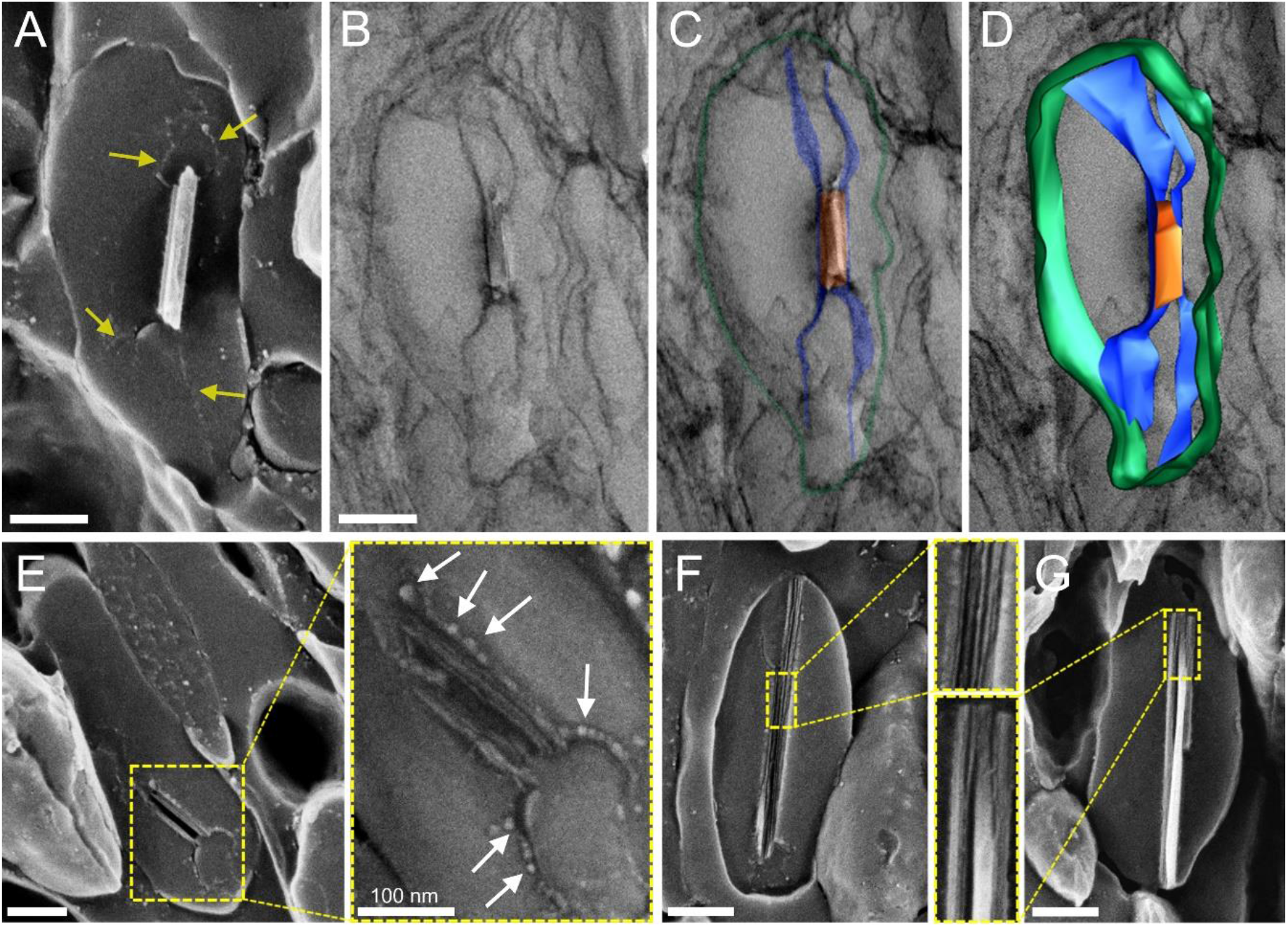
Intraluminal sheets control crystal nucleation and growth. **(A)** Cryo-SEM micrograph of an immature guanine crystal enveloped between two intraluminal fibrils (yellow arrows). **(B)** TEM micrograph of an iridosome containing an immature guanine crystal. **(C)** Pseudo-colored TEM of the same iridosome. Green; iridosome membrane, blue; intraluminal fibrils, orange; crystal. **(D)** Image segmentation obtained from 3D TEM tomography overlaid on the raw 2D TEM image, revealing that the intraluminal fibrils are 2D sheets. **(E)** Cryo-SEM image of a very immature crystal nucleating on the intraluminal sheets, which are decorated with tiny globules (white arrows). **(F)** Immature crystals are composed of 10-13 nm thick platelets which **(G)** coalesce to form thicker (25 nm) crystallites during maturation. Scale bars: 200 nm.

Cryo-SEM micrographs show that the intraluminal sheets are decorated with globular structures (Fig. 4E; white arrows). The immature crystals which nucleate on the sheets are constructed from 10-13 nm thick H-bonded platelets (Fig. 4E-F, Supplementary Fig. 5). In some cases, more than one nucleation point is observed (Supplementary Fig. 6). As crystallization proceeds, the platelets coalesce to form thicker (25 nm) crystallites (Fig. 3G). Similar merging of crystal platelets during guanine formation in spiders^17^ and lizards^29^, indicates that this may be a universal feature of guanine bio-crystallization.

Whilst ‘capping’ provides an answer to the formation of flat crystals, control over crystal twinning is the likely means of generating squares^26^. Scallop crystals are comprised of three monoclinic domains, twinned about the (100) plane. Evidence of this twinning is seen in the platelet texture of the crystals. Hirsch et. al., suggested^26^ that by controlling the number of domains and angle between twins, a crystal with a high symmetry morphology can be obtained from a low symmetry structure.

In addition to templating guanine nucleation and growth, the intraluminal sheets play a key role in controlling crystal orientation and organization. By mapping the orientation of sheets across different regions of the juvenile mirror (Fig. 5A-B), we determined that even prior to nucleation the sheets closely align with the curvature of the mirror. Thus, the directionality of the sheets in a juvenile scallop eye corresponds to the preferred orientation of the crystals in the adult obtained by *in situ* synchrotron XRD (Fig. 5C). This indicates that crystal orientation is ‘pre-programmed’ into the early iridosomes. Thus, the sheets are responsible for directing the highly reflective (100) crystal faces to the incident light and maintaining a smooth gradient of crystal orientations across the mirror surface - minimizing optical aberrations from surface defects^14^. Presumably, some ‘communication’ between organelles is required to generate the tessellated suprastructure, which is most probably achieved *via* the direction of cytoskeletal elements.

**Figure 5.**
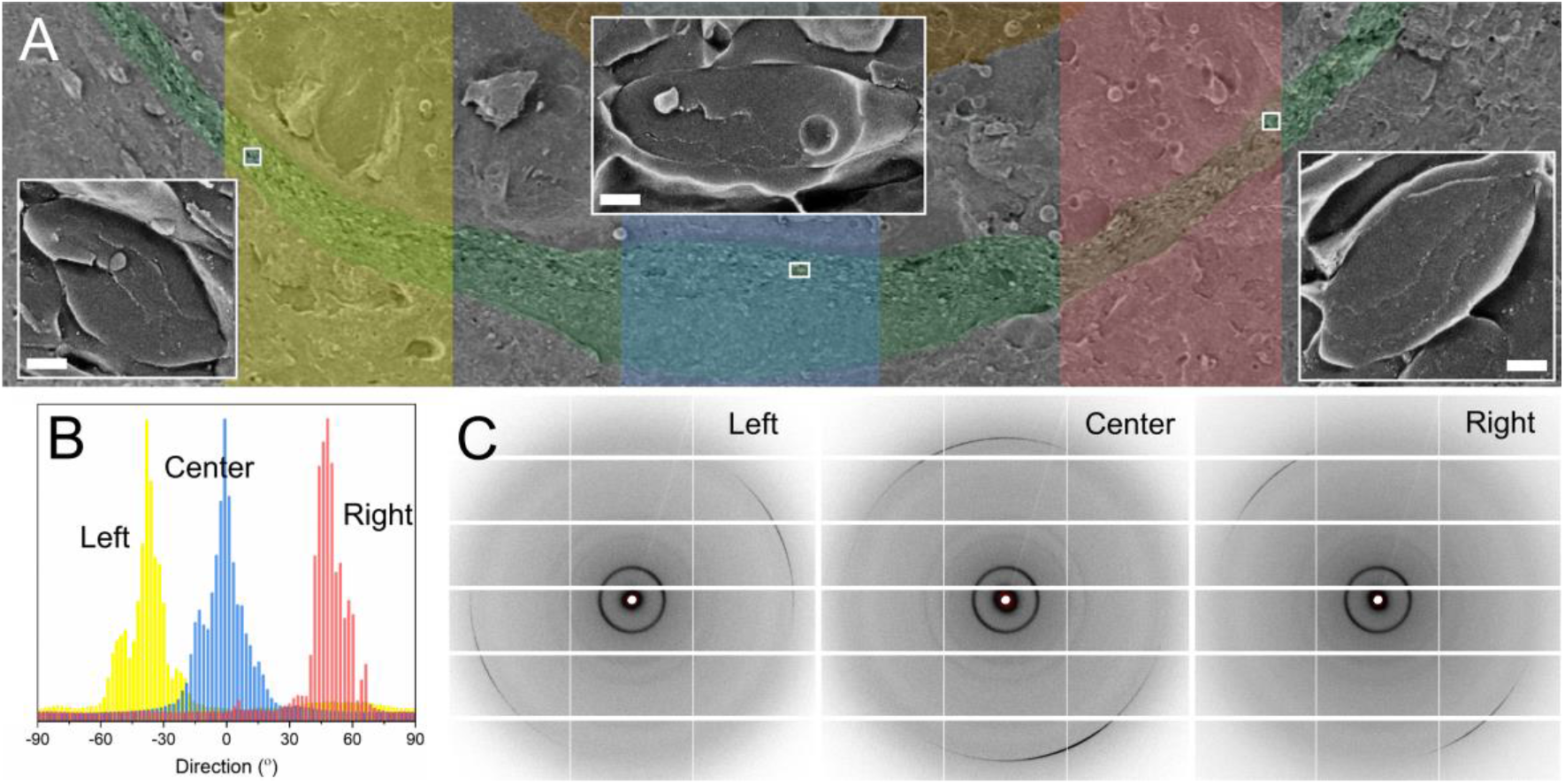
Intraluminal sheets control crystal orientation. **(A)** Low magnification cryo-SEM image of the mirror in a juvenile scallop eye (Fig.1B). Inserts: representative cryo-SEM images of iridosomes from the regions of interest, demonstrating the orientation of the sheets follows the curvature of the mirror. **(B)** Histograms representing the directionality of the templating sheets in the juvenile, from the three highlighted regions of interest in (B). **(C)** Representative *in situ μ*spot wide angle X-ray diffraction (WAXS) patterns from corresponding regions of the adult mirror showing the preferred orientation of the guanine (100) peak across the mirror. Scale bars: 200 nm.

## Discussion

By following the formation of guanine crystals in the image forming mirror of the scallop eye we have shown that pre-assembled macromolecular sheets within iridosome organelles template guanine crystal nucleation and growth. The role of these macromolecular sheets is twofold; they direct growth along the thermodynamically disfavoured H-bonding direction and orient the highly reflective face of guanine crystals towards focal points on the two retinas. These results provide an answer to the question of how organisms create highly reflective (100) guanine plates.

Another fundamental finding is the striking resemblance between the morphogenesis of guanine in scallops (Fig. 6; top row) and the well-documented melanosome organellogenesis in vertebrates (Fig. 6; bottom row)^30-32^. The four stages of melanosome formation in human MNT1 cells are shown alongside the key stages of guanine morphogenesis derived from Fig. 2. Melanosome formation begins with a spherical, endosomal compartment (a premelanosome) comprising small intraluminal vesicles that seed fibril formation^27^ (Fig. 6; bottom – stage I). The size and texture of these intraluminal vesicles and presence of disorganized fibrils are highly reminiscent of the multi-vesicular body of the early iridosome found here (Fig. 6; top – stage I, Fig. 2A). In stage II of melanosome formation (Fig. 6; bottom – stage II), amyloid sheets comprising the premelanosome protein (PMEL) assemble and stretch the organelle into an ellipsoid. As well as dictating melanosome shape, PMEL fibrils function to sequester highly reactive oxidative intermediates produced during melanin synthesis^33^. In iridosomes, prior to nucleation, exactly two highly oriented, intraluminal sheets assemble concomitantly with its transformation into an ellipse (Fig. 6; top – stage II). The 2D sheets then template the nucleation and growth of guanine (Fig. 6; top – stage III). This is analogous to stage III of melanosome formation, when tyrosine is oxidized to melanin, which is then deposited on pre-assembled PMEL sheets (Fig. 6; bottom – stage III). Though the chemistry of the templating sheets in iridosomes is unknown, the analogies to melanin suggest that they may be formed from a functional amyloid protein. Finally, following melanin and guanine deposition, the templating sheets remain an integral part of the organelle; being intercalated inside the pigment in melanosomes (Fig. 6; bottom – stage IV), and being fused to the guanine surface in iridosomes (Fig. 6; top – stage IV, S4).

**Figure 6.**
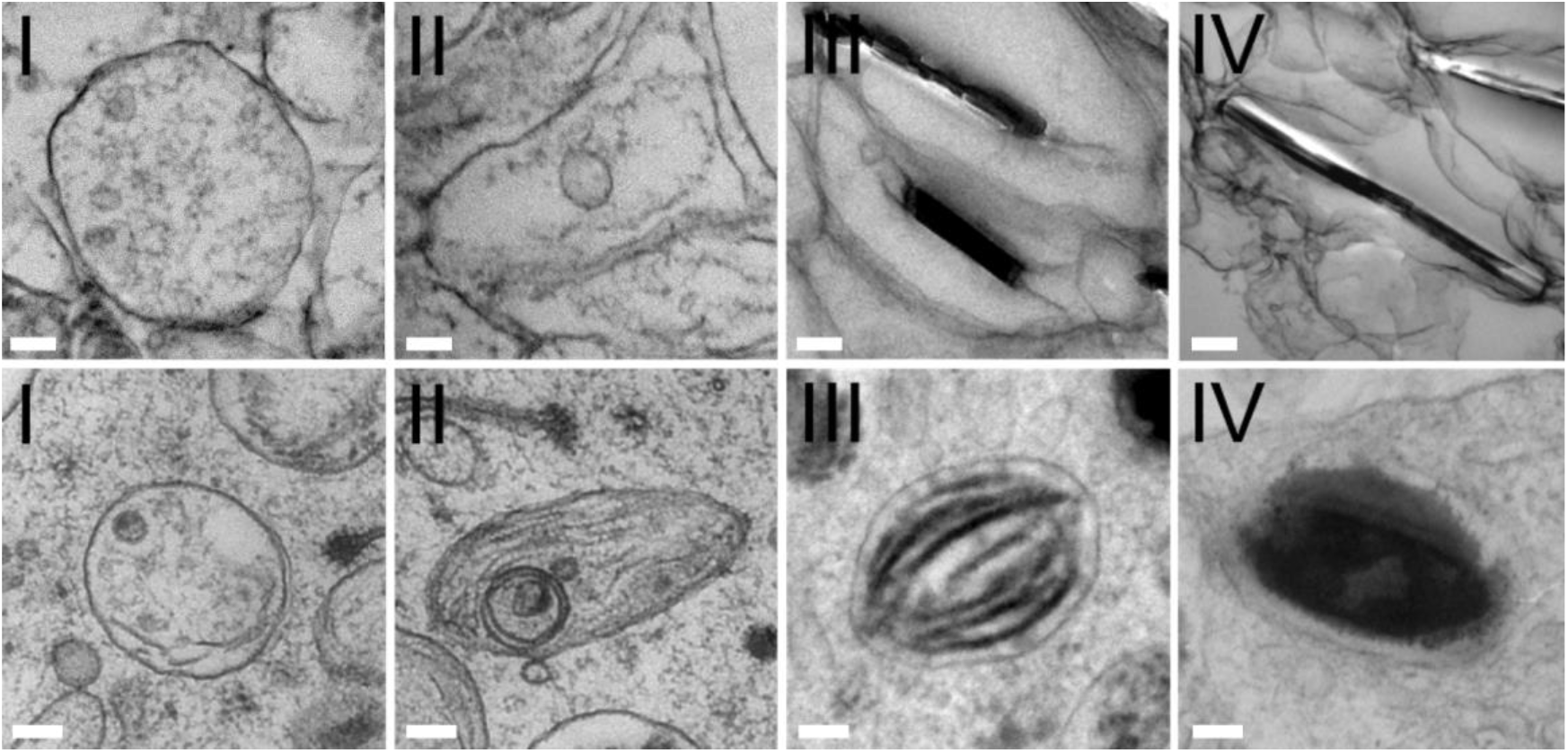
A comparison of iridosome and melanosome organellogenesis. **(Top)** TEM images of iridosomes from juvenile scallop eyes: Stage I iridosomes are spherical compartments containing ILVs and disorganized fibrils. Stage II iridosomes elongate concomitantly with the formation of two fibrillar sheets extending across the organelle. In stage III the sheets template the nucleation and growth of a guanine crystal and become bound to its (100) face. In stage IV, the growing guanine crystal stretches the organelle membrane tightly around itself. **(Bottom)** TEM images of melanosomes from high-pressure frozen MNT1 highly pigmented melanoma cells: Stage I melanosomes are spherical endosomal compartments containing ILVs which seed the formation of fibrils. These fibrils elongate and assemble into sheets of the PMEL protein, which stretch the organelle into an ellipsoid – stage II. The sheets template melanin deposition in stage III and IV. In stage IV melanosomes, the intercalated fibers are completely obscured by deposited melanin. Scale bars; 100 nm.

Due to their crystallinity and unique optical and physical properties, organic bio-crystals were, until now, usually treated within the framework of biomineralization not pigmentation. However, our results show that guanine crystal formation exhibits many parallels with the morphogenesis of other lysosome related pigment organelles. Specifically, we show that a universal control mechanism underlies guanine formation in scallops and melanin formation in vertebrates. In both iridosomes and melanosomes, pre-assembled intraluminal sheets are responsible for guanine/melanin nucleation and for dictating organelle shape. In iridosomes, the macromolecular sheets cap, and effectively inhibit growth along the favored π-stacking direction and promote growth along the H-bonding direction. This uncovers a long-sought explanation to how thermodynamically disfavored, high refractive index, plate-like biogenic guanine crystals are formed. Template-directed crystallization is utilized in many inorganic biominerals to control crystal orientation and properties^34,35^. However, its role in organic biocrystallization was not known hitherto. Identifying the chemistry of the intraluminal sheets and vesicles and determining how widespread this mechanism is in biology, will be key to the development of the field of organic biomineralization. Furthermore, understanding how macromolecular templates interact with biogenic organic crystals promises to yield new approaches for controlling the morphological properties of molecular materials^36^.

## Materials and Methods

### Specimen collection and preparation

Juvenile *Pecten maximus* scallops^37^ of different sizes were purchased from Scalpro AS. Scalpro AS produce scallop spat at their facilities at Rong, Øygarden County, Norway. Scallops from the local area were conditioned, spawned and larvae and spat raised to various sizes from settled larvae to 5-20 mm shell height. Spat are reared by Algetun AS and Hotate AS on the west coast of Norway to larger sizes (2-7 cm). With the help of Prof. Thorolf Magnesen from the University of Bergen, mantles were removed and preserved in a fixative solution of 4 % paraformaldehyde (PFA) and 2% glutaraldehyde (GA) in 1xPBS. In the case of very small scallops, less than 2 cm, the whole animal was fixed.

### Optical microscopy

A Zeiss Discovery.V20 stereomicroscope equipped with an Axiocam 305 color camera was used to take images of whole scallops, mantles, and eyes.

### Cryogenic Scanning Electron Microscopy (cryo-SEM) – sample preparation and imaging

Eyes from fixed scallops were sandwiched between two aluminum discs and cryo-immobilized in a high-pressure freezing (HPF) device (EM ICE, Leica). The frozen samples were then mounted on a holder under liquid nitrogen in a specialized loading station (EM VCM, Leica) and transferred under cryogenic conditions (EM VCT500, Leica) to a sample preparation freeze fracture device (EM ACE900, Leica). The samples were fractured at a speed of 160 mm/sec, etched for 5 minutes at -110°C, and coated with 3 nm of PtC. Samples were imaged in a HRSEM Gemini 300 SEM (Zeiss) by secondary electron in-lens detector while maintaining an operating temperature of -120°C.

### Preparation of ultra-thin tissue sections for TEM and STEM imaging of iridosomes

Chemically fixed scallop eyes were washed with 0.1M cacodylate buffer for 2 h and post fixed with 1% osmium tetroxide and 2% uranyl acetate according to the method published in Wagner et al.^17^. Ultra-thin sections were prepared with an ultra-microtome (RMC, Arizona, USA) and imaged with HRSEM Gemini 300 SEM (Zeiss) STEM detector, and Thermo Fisher Scientific (former FEI) Talos F200C transmission electron microscope operating at 200 kV. The images were taken with Ceta 16M CMOS camera. The electron diffraction patterns were obtained with a ThermoFisher Scientific (FEI) Tecnai T12 G^2^ TWIN TEM operating at 120 kV. Images and electron diffraction (ED) patterns were recorded using a Gatan 794 MultiScan CCD camera. ED analysis was done using Gatan Digital Micrograph software with DIFPack module. Images and ED patterns were recorded with consideration of potential beam damage to the sample, thus appropriate illumination conditions (spot size) were used to avoid it.

### High Pressure Freezing, sample preparation, and conventional EM imaging of melanosomes

MNT1, highly pigmented melanoma cells were seeded on carbonated sapphire discs and grown for 2 days. Cells were then immobilized by HPF using either a HPM 100 (Leica Microsystems, Germany) or a HPM Live μ (CryoCapCell, France), and freeze substituted in anhydrous acetone containing 1% OsO_4_/2% H_2_O for 64 h using the Automatic Freeze Substitution unit (AFS; Leica Microsystems). The samples were included in EMbed 812. Ultrathin sections were contrasted with an aqueous solution of uranyl acetate and Reynold’s lead citrate solution. Electron micrographs were acquired using a Transmission Electron Microscope (Tecnai Spirit; Thermo Fisher, Eindhoven, The Netherlands) operated at 80kV, and equipped with a 4k CCD camera (Quemesa, EMSIS, Muenster, Germany).

### TEM tomography: acquisition and analysis

TEM imaging and tomography were performed with Thermo Fisher Scientific (former FEI) Talos F200C transmission electron microscope operating at 200 kV. The images were taken with Ceta 16M CMOS camera. The tilt series were acquired with Thermo Fisher Scientific Tomography software (version 5.9) between angles varying from -60° to 60° with 1° steps. Before each acquisition of an image and before movement of the stage to the next tilt angle, a cross correlation of tracking and focusing were made. The post processing and the stack alignment were done with Thermo Fisher Scientific Inspect3D software (version 4.4). Later this was refined manually with the software Amira version 6.5 (Geometry Transforms/Align Slices). Movies were created on the movie creation panel of the animation director of Amira 6.5 (FEI, USA). Brightness and contrast levels were adjusted, and the movie was performed by animating the xy-ortho slice through the volume. Then it was exported with 25 frames per second, in the highest quality and as a MPEG file. Due to the low contrast between the cell wall, crystal, and filament, it was not possible to perform an automatic segmentation. Therefore, manual segmentation of the vesicle membrane, fibrils, and crystal was performed using IMOD 4.11 software^38,39^.

### Determining the long-range orientational ordering of the intraluminal sheets

Cryo-SEM images of different region of the mirror in a juvenile scallop eye were used for this analysis. The positions and orientations of the sheets were manually marked by representative lines. The resultant images underwent Fast Fourier Transformed (FFT) analysis using the Directionality plugin in FIJI imageJ (https://imagej.net/plugins/directionality)^40^. This plugin is used to infer the preferred orientation of structures in an image, which is visualized by a representative histogram.

### Synchrotron radiation in situ μspot wide angle X-ray diffraction

Wide-angle XRD (WAXD) measurements (Fig. 3J) were performed at the mySpot beamline, in the BESSY II synchrotron at Helmholtz-Zentrum Berlin für Materialien und Energie. Chemically fixed adult scallop eyes were mounted and sealed between (X-ray transparent) Kapton foil windows with a rectangular aluminum or lead frame. The samples were kept hydrated throughout the measurement with a drop of water inside the Kapton windows. As was described in detail previously^17^, an energy of 15 keV (λ = 0.826561 Å) was selected by a Mo/BC multilayer monochromator. The 2D WAXD patterns were measured using a Dectrix Eiger X 9M area detector (3000×3000 pixel, 75μm pixel size). A polycrystalline quartz standard was used to calibrate the beam center and the sample-detector distance. Visualization of the 2D scattering patterns was performed using DPDAK (Version 1.4.2). The data were normalized with respect to the primary beam monitor (ionization chamber) and corrected for background because of the pinhole and air scattering.

## Acknowledgments

We thank Profs. Steve Weiner, Lia Addadi and Michael S. Marks for stimulating discussions on this project. We acknowledge Alexey Tachlytski from Zeiss for his instruction regarding STEM operation and Tally Kossovsky from the Hebrew University of Jerusalem for her guidance and training on microtome operation. We acknowledge Lee Shelly for photographing the juvenile scallops (insert Fig 1A). We acknowledge Dr. Ivo Zizak and the BESSY II synchrotron, Helmholtz-Zentrum Berlin für Materialien und Energie, Germany, for provision of synchrotron time at the *μ*spot beamline. A.W. is grateful to the Azrieli Foundation for the award of an Azrieli Graduate Fellowship. B.A.P. is the Nahum Guzik Presidential Recruit and the recipient of the 2019 Azrieli Faculty Fellowship.

## Funding

This work was supported by an ERC Starting Grant (Grant number: 852948, ‘CRYSTALEYES’) awarded to B.A.P.

## Author contributions

Conceptualization: AW and BAP. Methodology: AW, AU and BAP. Investigation: AW, AU, RM, EZ, and GR. Resources: TM. Visualization: AW and RM. Funding acquisition: BAP. Supervision: BAP. Writing – original draft: AW and BAP. Writing – review & editing: AW, GR and BAP.

## Competing interests

The authors declare no competing interests.

## Data and materials availability

All the data that support the findings of this study are available from the corresponding author upon reasonable request.

## Notes

### Competing Interest Statement

The authors have declared no competing interest.

