## Supplementary material for "The Formation of Biogenic Guanine Crystals Closely Resembles Melanosome Morphogenesis": SI

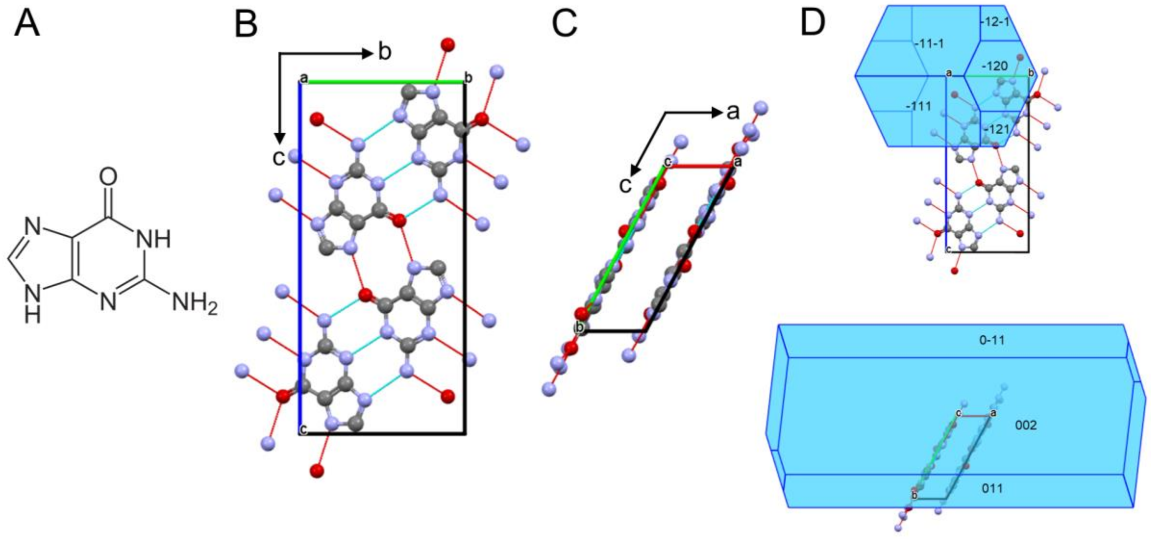


**Supplementary Fig. 1. The *β*-guanine crystal structure.** (A) Molecular structure of guanine. The crystal structure of *β*-guanine viewed perpendicular (B) and parallel (C) to the H-bonded layer[1]. (D) The corresponding thermodynamically stable BFDH morphology calculated by Mercury software.

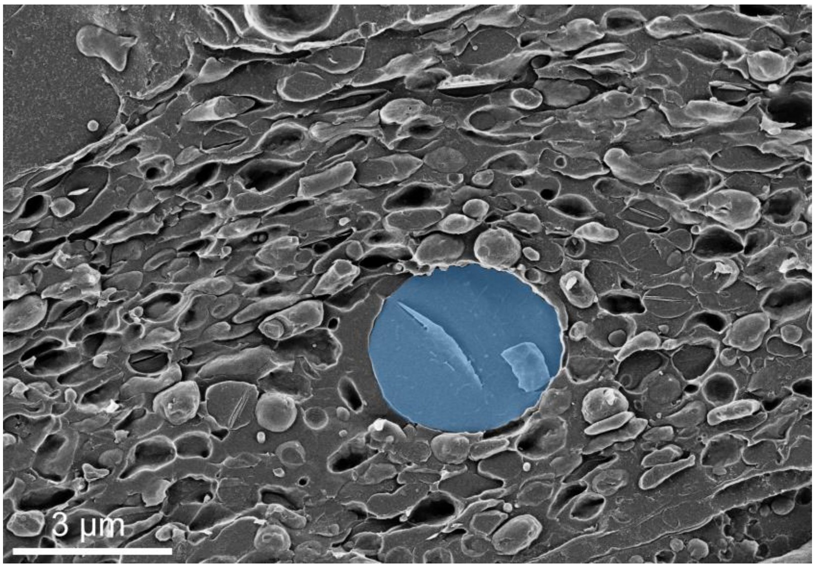


**Supplementary Fig. 2. Cryo-SEM micrograph of the mirror region in a freeze fractured juvenile scallop eye.** The nucleus (pseudo colored blue) of the iridophore is surrounded by ellipsoidal iridosomes of various stages, some containing partially formed guanine crystals.

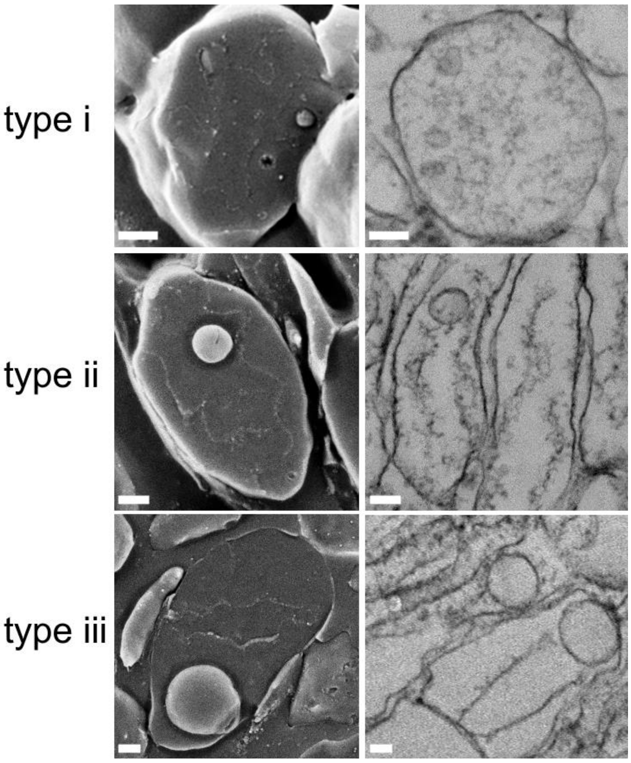


**Supplementary Fig. 3. The three different types of intraluminal vesicles (ILVs) observed in forming iridosomes.** Type *i* ILVs are between 25-50 nm in diameter and are associated with early spherical iridosomes and most probably have a role in fibril formation. Type *ii* ILVs are between 80-100 nm in diameter and are usually seen after fibril organization closely associating with one of the fibrils. Type *iii* ILVs are 250-350 nm in diameter and are seen after fibril formation either beside the fibrils or near one of the iridosome poles. Scale bars: 100 nm.

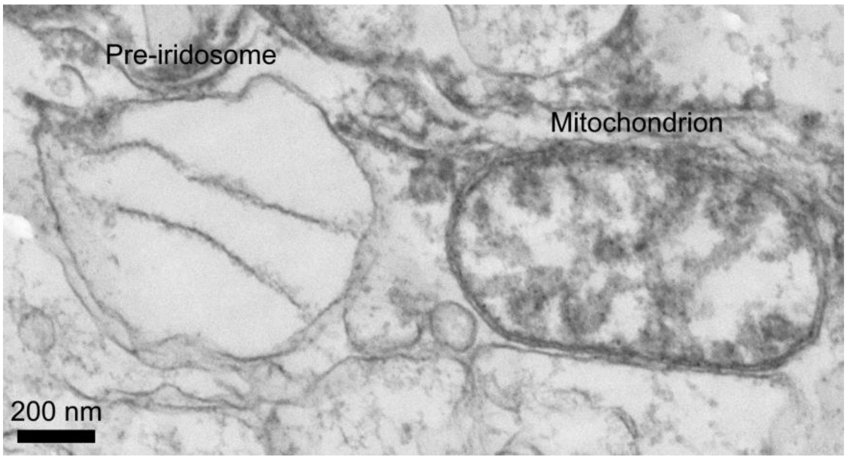


**Supplementary Fig. 4. The origin of "double-membrane" iridosomes.** Our observations not only describe the organellogenesis of iridosomes, but also clarify the origin of the "double-membrane" documented previously in these organelles[2-8]. The double-membrane was thought to derive either directly from the ER[4,7] or alternatively from fusion or incorporation of Golgi-derived vesicles with ER-derived vesicles[3,8]. Here we show the observed "inner membrane" is actually formed by the intraluminal sheets that template the crystal growth.

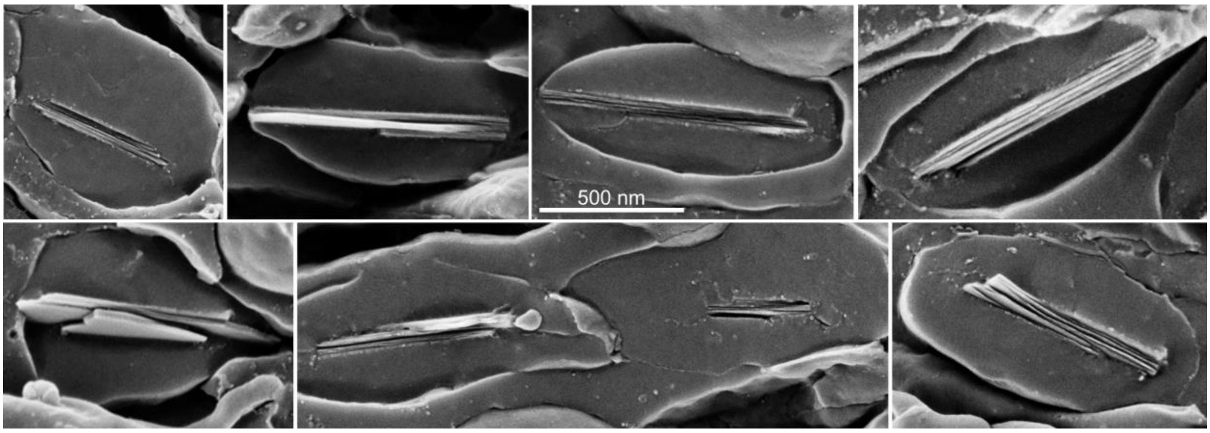


**Supplementary Fig. 5. Cryo-SEM images of immature crystals with layered texture.** The H-bonded guanine layers are oriented parallel to the (100) face of the crystal (parallel to the H-bonded plane). Initially each layer is 10-13 nm thick. As the crystal matures the layers coalesce to form thicker (25 nm) layers. Scale bar applies to all panels.

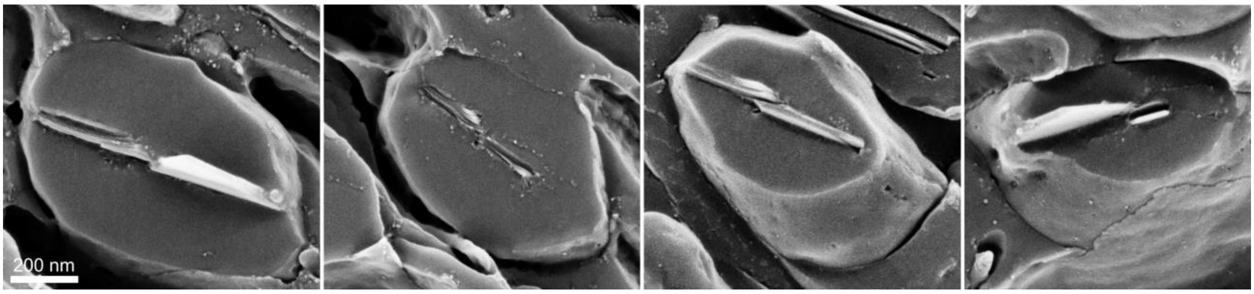


**Supplementary Fig. 6. Cryo-SEM images of crystals with multiple nucleation points.** The cryo-SEM images indicate that some guanine crystals have multiple nucleation points along the intraluminal sheets. Eventually the separately-nucleated crystals merge to form a single crystal. Scale bar applies to all panels**.**

**Supplementary Movie 1.**

TEM tomography movie of the juvenile scallop iridosome shown in Fig. 3B-D.

**Supplementary Movie 2.**

Additional TEM tomography movie showing a few iridosomes at different stages of development in a juvenile scallop.
